# Waves out of the Korean Peninsula and inter- and intra-species replacements in freshwater fishes in Japan

**DOI:** 10.1101/2020.10.05.325811

**Authors:** Shoji Taniguchi, Johanna Bertl, Andreas Futschik, Hirohisa Kishino, Toshio Okazaki

## Abstract

The Japanese archipelago is located at the periphery of the continent of Asia. Rivers in the Japanese archipelago, separated from the continent of Asia about 17 Ma, have experienced an intermittent exchange of freshwater fish taxa through a narrow land bridge generated by lowered sea level. As the Korean Peninsula and Japanese archipelago were not covered by an ice sheet during glacial periods, phylogeographical analyses in this region can trace the history of biota for a long time beyond the last glacial maximum. In this study, we analyzed the phylogeography of four freshwater fish taxa, *Hemibarbus longirostris*, dark chub *Nipponocypris temminckii, Tanakia* ssp. and *Carassius* ssp., whose distributions include both the Korean Peninsula and western Japan. We found for each taxon that a small component of diverse Korean clades of freshwater fishes migrated in waves into the Japanese archipelago to form the current phylogeographic structure of biota. Indigenous populations were replaced by succeeding migrants. We refer to this phenomenon as “waves out of the Korean Peninsula,” with parallels to “out of Africa” in humans.

## 1. INTRODUCTION

Inter- and intra-species interactions can influence biogeographical distributions (Pearson and Dawson, 2003; Waters, 2011; Wisz et al. 2013). Among many forms of biotic interactions, replacement among competing species that are mutually exclusive is presumed to be an important factor in biogeography (Gutiérrez et al., 2014; Yackulic, 2017). For humans, cultural records and genomic information have revealed a history of complex waves of dispersal and admixture out of Africa (Hellenthal et al., 2014; Nielsen et al., 2017).

While evidence for segregation is identified as genetic differentiation between geographical regions, phylogeographic evidence of intra-species replacements due to competition has not been extensively examined. Most evidence supporting the existence of species replacement is found in the spatial division and isolation of species, where the distributions of one species are surrounded by those of another (Gutiérrez et al., 2014). This contrasts with conventionally observed fragmentation, where populations are both genetically and spatially isolated from each other. Because fragmented populations are small, they have large genetic variation. They do not comprise a monophyletic group in phylogeny, but are interspersed by the main population. Conversely, local populations that have been recently divided by competitors of a different clade are genetically homogeneous and comprise a monophyletic group (Figure 1).

**FIGURE 1.**
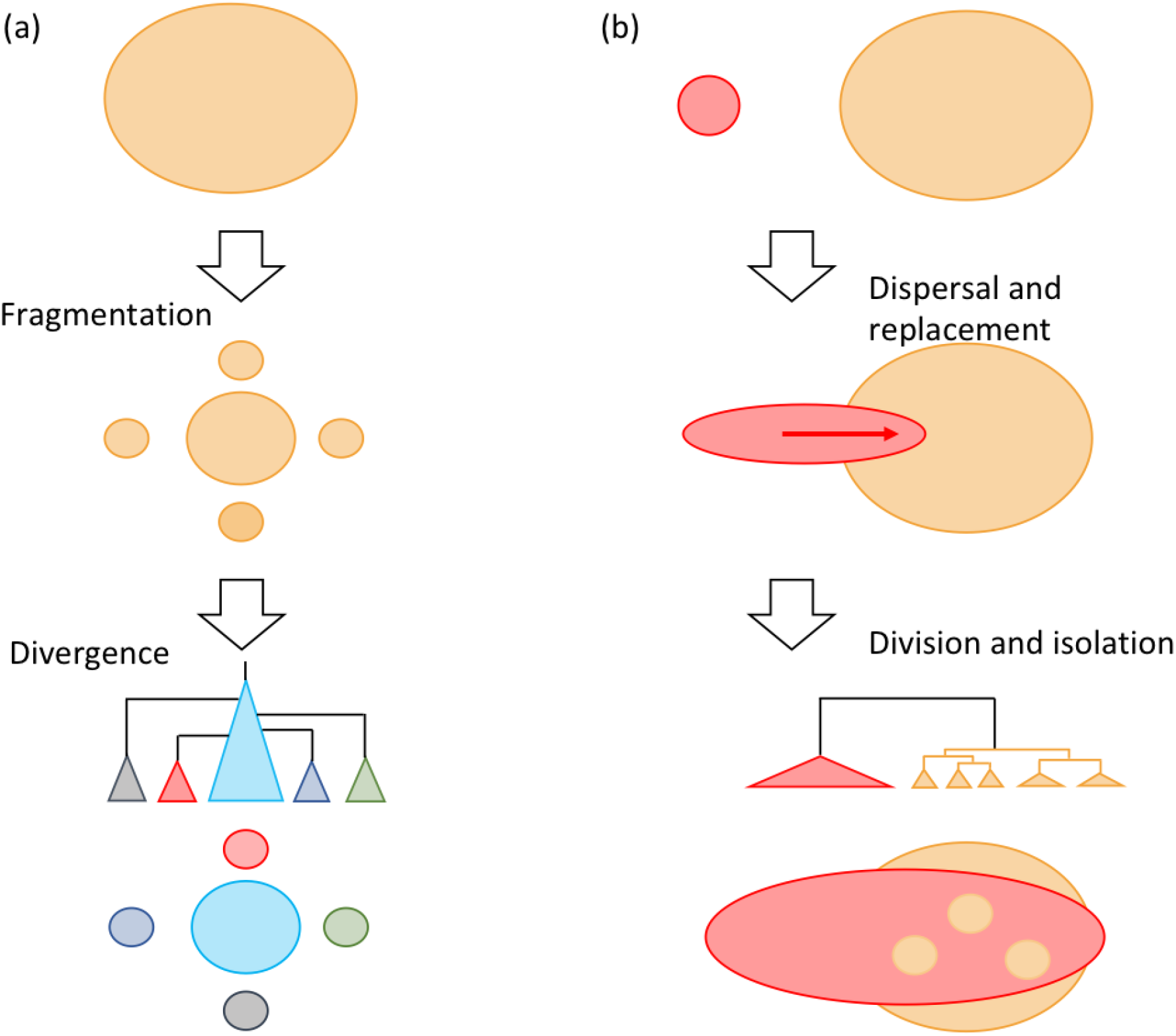
Schematic illustration of conventional fragmentation and replacement. (a) conventional fragmentation, where fragmented populations diverge into small populations with unique genetic features. Isolated populations do not comprise monophyletic groups in the phylogeny, but are interspersed by the main population. (b) Local populations (three small circles and upper and lower fragments) recently divided by competitors from different clades are genetically homogeneous and comprise monophyletic groups.

Gene flow among freshwater fishes is extremely low, as salinity barriers separate neighboring rivers. Dispersal within freshwater systems is prevalent, while dispersal between them is rare, with watersheds and oceans representing migration barriers. Between-system dispersal can occur by floods in lower basins, stream capture, or the appearance of a larger freshwater system that connects rivers in periods of decreased sea level during glacial periods. Limited genetic exchange between rivers lowers rates of species replacement. In the Quaternary, the distribution of biota was influenced by ice-sheets (Shafer et al., 2010). In northern hemisphere mid-latitudes such as throughout Europe or North America it is difficult to trace biogeographic history beyond the most recent ice-sheet formation of about 10 Ka. However, the Japanese archipelago and Korean Peninsula were never covered by ice sheets during glacial periods (Flint, 1971). Therefore, many taxa suitable for appraising the effects of competition on the distributions of genetic clades might exist in this region.

The Japanese archipelago is located at the periphery of the continent of Asia. The archipelago landmass originally formed the eastern margin of the continent of Asia. After the back-arc of the archipelago opened about 17 Ma, the northeastern half rotated counter-clockwise, while the southwestern half rotated clockwise. The current position was reached about 14 Ma (Baba et al., 2007), and fused into the current Japanese archipelago about 6 Ma. The boundary between these northeastern and southwestern masses is called the Fossa Magna (Figure 2a). The Japanese archipelago is elongated in a bow along a north–southwest axis. The mountains extending along this archipelago generate numerous short rivers that discharge separately into the ocean. The Sea of Japan is deep and has isolated the islands from the continent of Asia, except for narrow bridges at either end during periods of lowered sea level. These access points provide potential routes for genetic exchange of freshwater fishes on the Korean Peninsula with those on the Japanese archipelago. In the Japanese archipelago, the bottom of the Inland Sea was above sea level in glacial periods, and paleo-river systems connecting surrounding rivers (Figure 2b) enabled gene flow (Watanabe et al., 2017). Therefore, the current Inland Sea probably represented a likely dispersal route. Eastern dispersal was blocked after the uplift of Suzuka and Nunobiki Mountains at about 1 to 1.5 Ma.

**FIGURE 2.**
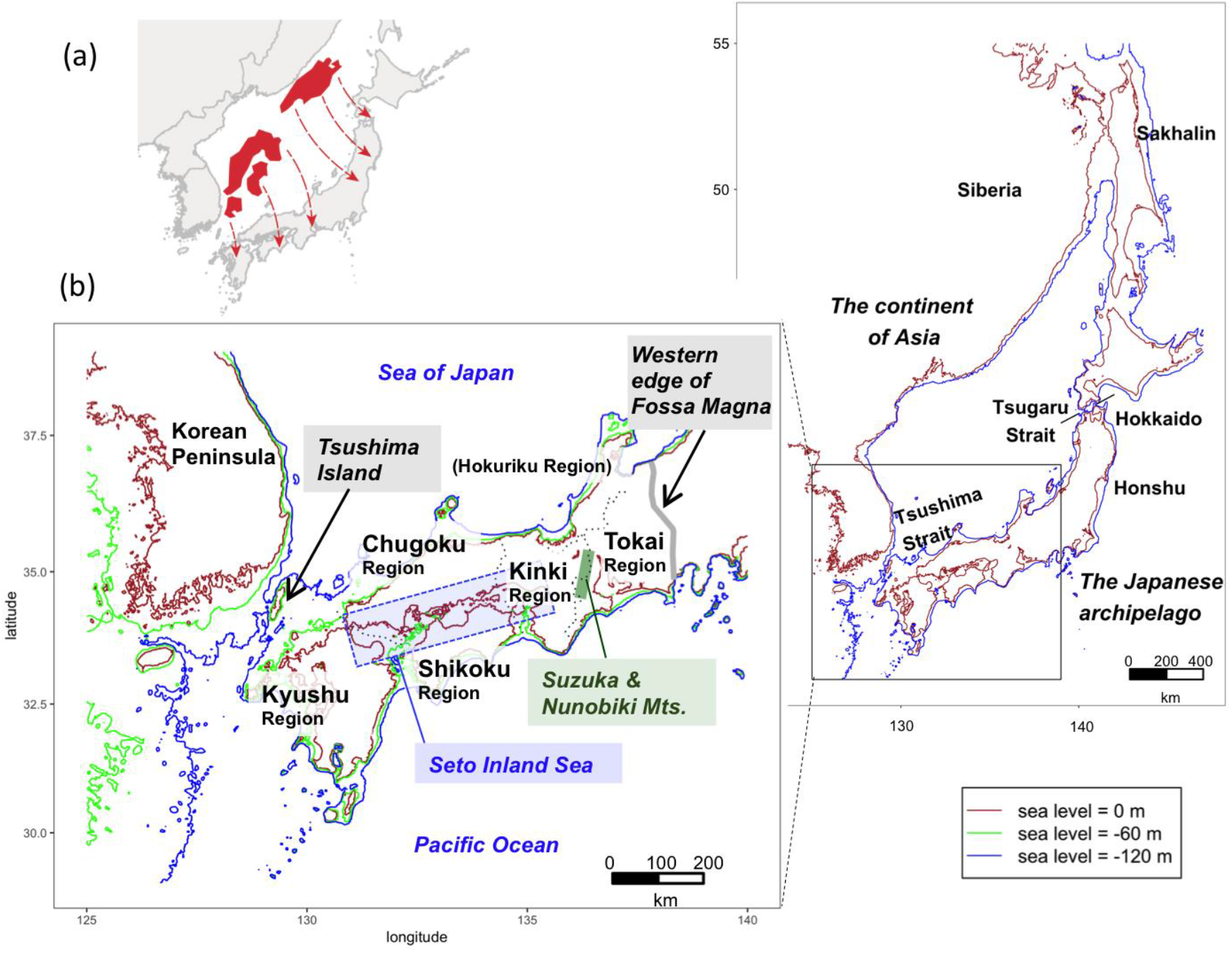
History of the Japanese Archipelago and study area. (a) Paleogeography of the Japanese archipelago inferred from geophysical information (Otofuji, 1996). (b) Study area map. An inferred map of the Far East during the last glacial period. The Seto Inland Sea, surrounded by Kyushu, Chugoku, Shikoku, and Kinki regions, with an average water depth of 38 m. Green and blue lines represent 60 m and 120 m bottom depths (Fairbanks, 1989; Rohling et al., 1998), respectively, and allow inferences to be made about coastal lines at times of lowered sea level.

A key factor in the establishment of the freshwater fish fauna of Japan is the isolation of these islands and their rivers, which affected the rates of expansion of migrant faunas. Here, we examine the effects of the migration and intra- and inter-species replacements on the phylogeographic structure in Japan. For this purpose, we analyze the phylogeography of four freshwater fish taxa, *Hemibarbus longirostris* (Regan, 1908), dark chub *Nipponocypris temminckii* (Temminck & Schlegel, 1846), *Tanakia* ssp. and *Carassius* ssp., whose distribution includes the Korean Peninsula and western Japan. The first three taxa have never been the target of commercial fisheries. Phylogenetic analysis combined with the geographical distributions of clades reveals that migrants from the Korean Peninsula have replaced indigenous populations in Japan. A simulation-based Bayesian analysis of *N. temminckii* reveals the former dominated the latter significantly.

For humans, it has been reported that series of waves of man originated in Africa and propagated around the world. When these waves interacted with pre-existing populations, hybridizations and sometimes replacements occurred along the way in a phenomenon known as “out of Africa.” In East Asia, a component of diverse populations of freshwater fishes in Korea migrated in waves into the Japanese archipelago; intra- and inter-specific competition between these new migrants and pre-existing populations then resulted in the current phylogeographic structure of Japanese freshwater fish biota.

## 2. Materials and Methods

### 2.1. Sampling and sequencing

Over 30 years, Toshio Okazaki (T.O.) amassed samples of freshwater fish from throughout Japan and Korea. From each sampling site, representative fishes were collected by various procedures, such as netting and angling. Fishes were fixed appropriately for molecular analyses [ETOH or −20°C]).

#### Hemibarbus longirostris

samples were collected between 1990 and 2009 in Japan, and between 1991 and 1994 in South Korea. Among them, 27 individuals from 15 sites in Japan and 63 individuals from 20 sites in South Korea were subjected to analysis. By conducting PCR-RFLP analysis with 15 × 4-base pair recognition restriction enzymes (Appendix A, supplementary information), individuals were selected for sequencing. At each site, one individual was selected for sequencing when all individuals from a site had the same banding pattern. All individuals with different banding patterns were subjected for sequencing. To extract high resolution phylogeographic information, we chose the rapidly evolving mitochondrial gene NADH dehydrogenase subunit 2 (ND2) as a molecular marker. As a result, 42 × ND2 sequences of 584bp were obtained. We sequenced the same rapidly evolving ND2 for other freshwater fishes.

#### Nipponocypris temminckii

samples were collected between 1988 and 2013 in Japan, and between 1991 and 1994 in South Korea. Among them, 561 individuals from 340 sites in Japan and 93 individuals from 57 sites in South Korea were subjected to analysis. By conducting PCR-RFLP analysis with 13 × 4-base pair recognition restriction enzymes (Appendix A), individuals were selected for sequencing. At each site, one individual was selected for sequencing when all individuals from a site had the same banding pattern. Also, when individuals from many geographically close streams had the same banding pattern, individuals from equidistant streams were subsampled for sequencing. All individuals with different banding patterns were subjected for sequencing. As a result, 309 individuals from 248 sites in Japan and 41 individuals from 32 sites in South Korea were sequenced. We sequenced a partial region of ND2, 600 bp. To calculate the rate of ND2 molecular evolution, cytochrome b sequence data were obtained from 22 samples of individuals from all clades. Sequencing followed procedures described in Appendix A (supplementary information).

Oily bitterling *Tanakia limbata* (Temminck & Schlegel, 1846), *T. koreensis* (Kim & Kim, 1990) and related species: samples were collected between 1989 and 2013 in Japan. Among them, 97 individuals from 47 sites were subjected to analysis. Samples of *T. koreensis* and related species were collected between 1991 and 1994 in South Korea. Among them, 17 individuals from 15 sites were subjected to analysis. As for genus *Tanakia*, individuals were screened for sequencing based on the PCR-RFLP analysis with 15 × 4-base pair recognition restriction enzymes (Appendix A). As a result, 70 × ND2 sequences of 741bp were obtained.

*Carassius auratus* (Linnaeus, 1758), *C.* sp. and Japanese white crucian carp *C. cuvieri* Temminck & Schlegel, 1846: samples were collected between 1989 and 2007 in Japan, and between 1991 and 1994 in South Korea. Among them, 274 individuals from 48 sites in Japan, and 101 individuals from 41 sites in South Korea were subjected to analysis. Three goldfish that we analyzed have been exploited commercially, unlike *N. temminckii* and *T. limbata*. Largely because of between-river transplantations, the PCR-RFLP analysis with 10 × 4-base pair recognition restriction enzymes (supplementary information) did not identify haplotypes that characterized regions. Therefore, all individuals with different banding patterns were sequenced. As a result, 154 × ND2 sequences of 600bp were obtained. The obtained sequences were deposited in DDBJ/ENA/GenBank (accession numbers were LC566827-LC566870, LC566893-LC567047, LC567225-LC567298 and LC567958-LC568289).

### 2.2. Sampling sites and geomorphological information

Precise collection sites were identified by T. O. based on field notes. Bathymetry was determined from ETOPO1 data (Amante and Eakins, 2009) using R (R Core Team, 2017) marmap (Pante and Simon-Bouhet, 2013). A shaded-relief map was made from the elevation chart (Geospatial Information Authority of Japan, 2013), with marine areas assembled using data from the Hydrographic and Oceanographic Department, Japan Coast Guard. River data were obtained from the National Land Numerical Information download service (Ministry of Land, Infrastructure, Transport and Tourism, 2011).

### 2.3 Phylogenetic analysis: *H. longirostris*

A phylogenetic tree was constructed using the maximum likelihood (ML) method implemented in MEGA7 (Kumar et al., 2016). We used sequences of *H. mylodon* (Berg, 1907) and barbel steed *H. barbus* (Pallas, 1776) as outgroups for *H. longirostris*. The Bayesian Information Criterion (BIC) selected the TN93+I model as the best model of nucleotide substitution. Bootstrap values of the ML-estimated branches were obtained by analyzing 1,000 resampled alignment columns. Based on the sequence-based phylogenetic tree and sample sites, we reconstructed ancestral distributions of species throughout Japan and Korea using ML (Pagel, 1994). To account for uncertainty in the phylogenetic tree, we constructed our tree using the Bayesian MCMC method implemented in BEAST 2.4.7 (Bouckaert et al., 2014). We extracted a 10% subsample of the MCMC sample of 9,000 trees by sampling at regular intervals and used it as an input together with trait data for ML analysis by BAYESTRAITS V3. We confirmed that the phylogenetic clades of the current *H. longirostris* population in Japan were formed by a migration event from Korea.

### 2.4. Phylogenetic analyses: *N. temminckii*

The 309 sequences for *N. temminckii* included 109 unique sequences. A phylogenetic tree was constructed using the maximum likelihood (ML) method implemented in MEGA7 (Kumar et al., 2016), and the Bayesian method implemented in BEAST 2.4.7 (Bouckaert et al., 2014); *N. sieboldii* (Temminck & Schlegel, 1846) was used as an outgroup. BIC selected the TN93+I model as the best model of nucleotide substitution. In the Bayesian tree inference, we assumed TN93+I as the model of nucleotide substitution. The coalescent process model of a population in equilibrium was adopted as a prior for the tree. As a prior on the rate of molecular evolution, we assumed the log-transformed values followed an autocorrelated normal distribution (Drummond et al., 2006). The mean evolutionary rate was obtained by converting the frequently cited cytochrome b molecular evolutionary rate, 0.76% per site per MY (Zardoya and Doadrio (1999)), to that of ND2. By comparing the average evolutionary distance of cytochrome b sequences (0.0627) and corresponding ND2 sequences (0.0513), the ND2 evolutionary rate was estimated at 0.93%. We estimated divergence times in the Bayesian framework using the MCMCTREE package implemented in PAML 4.9 (Yang, 2007). Detailed methods are described in Appendix A1.

#### 2.4.1 Geographical origin

Based on the sequence-based phylogenetic tree and sample sites, we reconstructed ancestral distributions of species throughout Japan and Korea by the same method as for *H. longirostris*. We confirmed that the phylogenetic clades of the current *N. temminckii* population in Japan were formed by multiple migration events from Korea.

#### 2.4.2. Simulation-based testing of the intra-species replacements

To test our hypothesis that indigenous populations of fishes were replaced by new migrants, we simulated the formation process for *N. temminckii*. By contrasting simulated patterns in the distribution of this species under various competitive scenarios with the observed pattern, we examined if information contained in sequences and their sites provided evidence for intra-species replacement. Simulation assumes the timing of migration from Korea, dispersal rate within the Japanese archipelago, and replacement rate among intra-species clades, specifies the formation process. By estimating these parameters via approximate Bayesian computation (ABC; Beaumont et al., 2002), the significance of the biogeographic evidence on intra-species replacement was tested.

As summary statistics sensitive to model parameters (Aeschbacher et al., 2012), we extracted features from the geographical distributions of clades that contained information on the timing of migration, and dispersal and replacement rates. These Templeton statistics were used in Nested Clade Phylogeographic Analysis (NCPA; Posada et al., 2006; Templeton et al., 1995) and spatial autocorrelation (Figure B1). NCPA was designed to associate the haplotype tree and geography through clade and nested clade distances, and to infer evolutionary history from population structure (Templeton, 2004). Spatial autocorrelation measures the correlation between genetic and geological distances and has a small value when a sample includes individuals that are surrounded by different neighboring clades. For a detailed explanation of the simulation algorithm, see Appendix A1.

### 2.5. The case of *T. limbata* and *Carassius* and their related species

ML phylogenetic trees and biogeographic maps for *T limbata, Carassius* and their related species were obtained in a similar fashion to those for *N. temminckii*. BIC selected the TN93+G+I model as the best model of nucleotide substitution of *T. limbata* and TN93+I for genus *Carassius* respectively. We used sequences of *T. lanceolate* (Temminck & Schlegel, 1846), Tokyo bitterling *T. tanago* (Tanaka, 1909), *Acheilognathus rhombeus* (Temminck & Schlegel, 1846), and big-scaled redfin *Tribolodon hakonensis* (Günther, 1877) as outgroups for *T. limbata*, and *Cyprinus carpio* Linnaeus, 1758 as the outgroup for *Carassius*.

### 2.6. Data availability

Sequence data and sampling sites are available at NCBI. All freshwater fishes sampled by T.O., including *H. longirostris, N. temminckii, C. cuvieri* and *T. limbata*, are kept at Seikai National Fisheries Research Institute. Dr. Koichi Hoshino is responsible for the “Okazaki collection.”

### 2.7. Code availability

All analysis scripts can be found at https://github.com/ShojiTaniguchi/Division_and_Isolation.

### 2.8 Ethical Statement

The guidelines or policies on conducting animal experiment are stated in “Act on Welfare and Management of Animals” enacted in 1973 revised in 1999 and “Standards relating to the Care and Keeping and Reducing Pain of Laboratory Animals” (Notice of the Ministry of the Environment No. 88 of 2006). Most of the specimens used for the analysis were collected before 2000 while T.O. was working for National Research Institute of Aquaculture and National Research Institute of Fisheries Science, which also set up its guideline on animal experiment in 2008. However, fish is not subject to any of the above regulations or the guideline. Since there were no regulations or guidelines regarding the handling of fish body, the fish specimens were either immediately frozen or preserved in alcohol. The fish species we collected are commonly distributed and certainly not designated as nationally protected species or subject to any regulations.

## 3. Results

### 3.1 Phylogeography of *H. longirostris*

ML mt tree for *H. longirostris* identified three monophyletic clades (I, II, III). All individuals in clades I and III were sampled in South Korea, while clade II included individuals from South Korea and Japan (Figure 3). Clade I is distributed in the central region of the Korean Peninsula, and Clade III is distributed in the southeastern region of Korea. The distribution of clade II includes two isolated areas across the Tsushima Strait—the southeast region of Korea, and western Japan, the latter distribution expanding from the Chugoku region facing the Seto Inland Sea (hereafter ‘Inland Sea’) to the western part of the Tokai region. Clade II originated in Korea, with a main migration event into Japan. This indicates the three clades diverged in Korea, with a clade splitting from the Korean population and migrating onto the Japanese archipelago via a land bridge.

**FIGURE 3.**
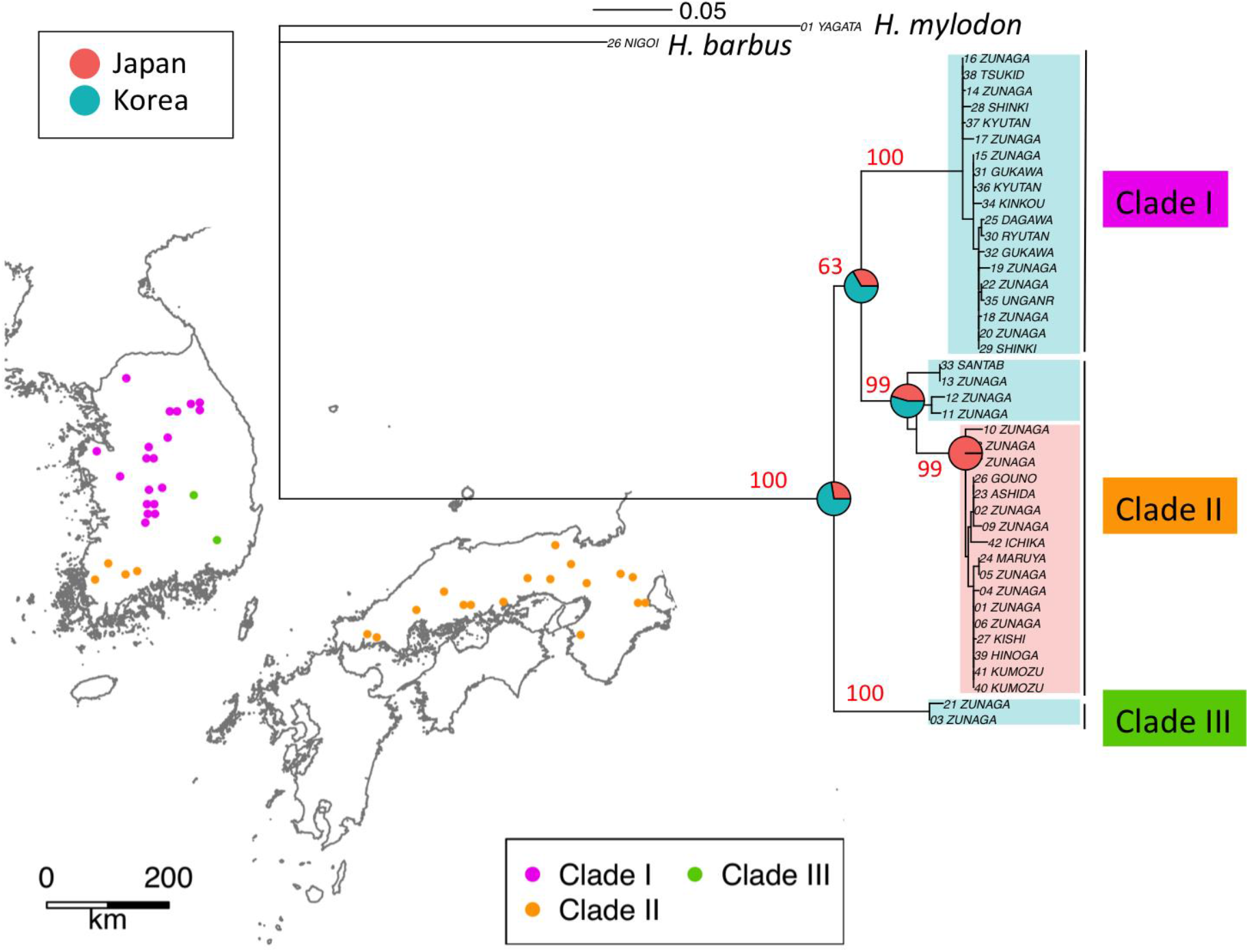
Phylogenetic trees of ND2 sequences and geographic locations of *Hemibarbus longirostris* reconstructed from ND2 sequences. Red numbers in the phylogenetic tree indicate bootstrap probabilities (%). Pie charts on the phylogenetic tree represents the Bayesian assignment of ancestral nodes to Japan and Korea.

Two land bridges between the continent and Japan are considered possible migration routes: the first between Korea and western Japan, and the second between the maritime province of Siberia and Hokkaido in Japan, through to Sakhalin in Russia (Figure 2b). The deep Tsugaru Strait between Honshu (the main island) and Hokkaido represented a dispersal barrier for freshwater fishes (Watanabe, 2012). Since their natural habitat does not extend to eastern and northern Japan, *H. longirostris* apparently used the land bridge to migrate from Korea to western Japan.

### 3.2. Phylogeography of *N. temminckii*

ML and Bayesian mt trees for *N. temminckii* consistently identified seven clades (A–G, Figure 4a, B2a, B2b). Monophyly of clade F was significant in the Bayesian tree. All individuals in clades A, B, and D were sampled in South Korea, while clade C included individuals from South Korea and Japan (Figure 4b). The Korean population (C) is currently regarded as a distinct species, *N. koreanus* (Kim, Oh & Hosoya, 2005). Individuals in clade F were sampled in Japan, while those in clade G included individuals from both Japan and Korea. Clade F is distributed in the regions of Kinki, Chugoku, and Shikoku, whereas clade G is distributed in the regions of Chugoku, Shikoku, and Kyushu (Figure 4b). The distributions of the two clades appear to overlap in Chugoku and Shikoku, but fine-scale sampling in these regions reveals they are actually segregated (Figure 4c).

**FIGURE 4.**
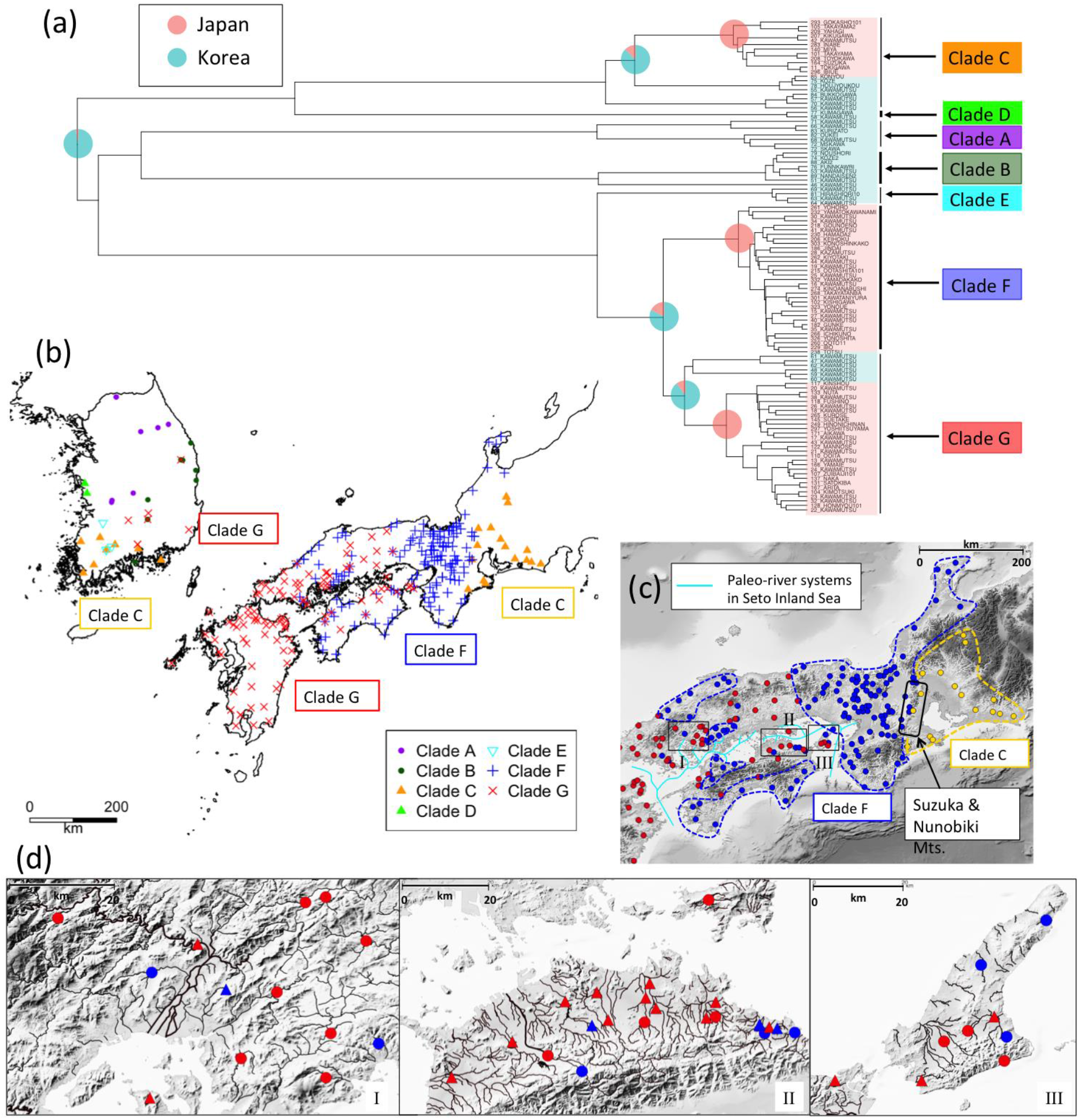
Phylogeny and distribution of *Nipponocypris temminckii*. (a) Bayesian assignment of ancestral nodes to Japan and Korea; (b) sample sites by clade; (c) distribution of clades on shaded-relief map; and (d) focused distributions at sites I, II and III. Detailed distributions of individuals for which the type of a clade has been determined by PCR-RFLP analysis are displayed as triangular points (d). Colors represent the cleavage types of BstUI, DdeI, and TaqI that correspond to clades F and G (see supplementary information). Paleo-river systems (c) modified from original maps (Japan Association for Quaternary Research, 1987; Kuwashiro, 1959). At localities I and III, clade F was sampled at sites surrounded by mountains, but at locality II, clade F was sampled upstream and clade G downstream. We used R (R Core Team, 2017) package GGTREE (Yuet al., 2017) for visualization in (a).

The distribution of *N. temminckii* in clade C includes two largely separated geographical regions—the southeast region of the Korean Peninsula, and the Tokai region of Japan. Clades F and G occupied areas west of the Tokai region (Okazaki et al., 1991). Clade G is widely distributed throughout the southern region of Korea, and Kyushu, Chugoku, and Shikoku in Japan, whereas clade F is absent in Kyushu, but mostly present in Kinki (Figure 4b, 4c). A few samples of clade F were obtained in Tokai, at the upper reaches of the rivers running eastern side of Suzuka Mountains. Clade G expands around the Inland Sea, as does clade F, which is also found on the Sea of Japan side of Chugoku, and the Pacific Ocean side of Shikoku (Figure 4c), both isolated from the Inland Sea by mountains. In some areas, fishes in clade F occurred upstream or in mountain locations, while those in clade G occurred in the same river system in the mainstream or downstream (Figure 4d). The three clades originated in Korea, with three main migration events into Japan (Figure 4a). Clades C, F and G diverged in Korea, with some clades splitting from the Korean population and migrating onto the Japanese archipelago via a land bridge. For reasons similar to *H. longirostris, N. temminckii* apparently used the land bridge to migrate from Korea into western Japan.

Since clade C inhabits the Tokai region, an area farthest from the land bridge in the Japanese distribution of *N. temminckii*, it represents the oldest clade in Japan (Figure 4b). Similarly, clade F is likely to be the second oldest clade, as it is distributed in the area second farthest from the land bridge. Clade G represents the youngest clade, as its distribution throughout Japan and Korea is closest. The existence of the Fossa Magna east of Tokai and Hokuriku (Figure 2b) might explain why *N. temminckii* did not expand further east. In glacial periods two paleo-river systems in the Inland Sea (Figure 4c) enabled gene flow (Watanabe et al., 2017). Therefore, the current Inland Sea was a likely dispersal route. The consecutive migration of clades C, F and G, followed by their dispersal might explain the present distribution of this species in Japan. Estimated divergence times are consistent with this scenario. As shown in Figure B3, node 1 (the split of clade C between Korea and Japan) is older than node 2 (the period of migration of clade F), with a posterior probability of 83.8%. Additionally, node 1 is older than node 3 (the period of migration of clade F from Korea), with a posterior probability of 98.2%. Although the phylogenetic tree does not provide a decisive chronology of nodal migration, the distribution map clearly suggests node 1 is older (Figure 4b). Bayesian tree estimation (Figure B2b) reveals that migration of clade C occurred 1.52 Ma [0.876–2.04], clade F 1.31 Ma [0.896–1.74], and clade G 1.12 Ma [0.713–1.43]. Therefore, we assume that the split in clade C between Korea and Japan occurred first, with clade F and G diverging from a common ancestor.

### 3.3 Significance of intra-species replacement

The *N. temminckii* clades F and G are similar, with average nucleotide distances of 1.78%, far lower than genetic distances among salmonid species (Thomas & Beckenbach, 1989). The taxonomic status of clade C is confusing; its population in Japan is regarded as the same species as clades F and G, while the population in Korea is considered to represent *T. koreensis*. The divided pattern of clade C is presumably at the intra-species level while the taxonomy requires review. Distributions of *N. temminckii* within Japan were simulated using various parameters (Figure 5a). The initial state was a universal distribution of clade C in western Japan. Clades F and G migrated into western Japan and expanded their distributions. Depending on replacement rate, resultant distributions differed (Figures 5b, 5c). To avoid effects of differing sampling effort, we considered dispersal of individuals on a lattice-like grid in Japan. Our model assumes dispersal distance follows a gamma distribution; it has four parameters: r (Ma) for the timing of migration of clade G, *m* for the dispersal rate (km/Kyr), *s* (km) for the scale parameter of the gamma distribution, and *α* for replacement rate. The timing of migration of clade F was set to 1.31 Ma, inferred using a Bayesian procedure (Figure B2b). Under this model, *α* was estimated at 0.774 [0.554–0.951] (Figure B4), which is significantly higher than that for a neutral relationship (0.5). We therefore reject the null hypothesis of selective neutrality that assumes that the three clades C, F, and G have equal fitness. Of other parameters, *m* was extremely low, estimated at 0.345 [0.0135–0.860] km/Kyr, and *s* at 20.2 [5.33–40.1] km. The point estimates of *m* and *s* indicate that short dispersal was more frequent than long dispersal. The migration of clade G was dated at 0.862 Ma [0.552–1.30]. The MCMCTREE gave a conditional credibility interval of 0.862 Ma [0.543–1.238].

**FIGURE 5.**
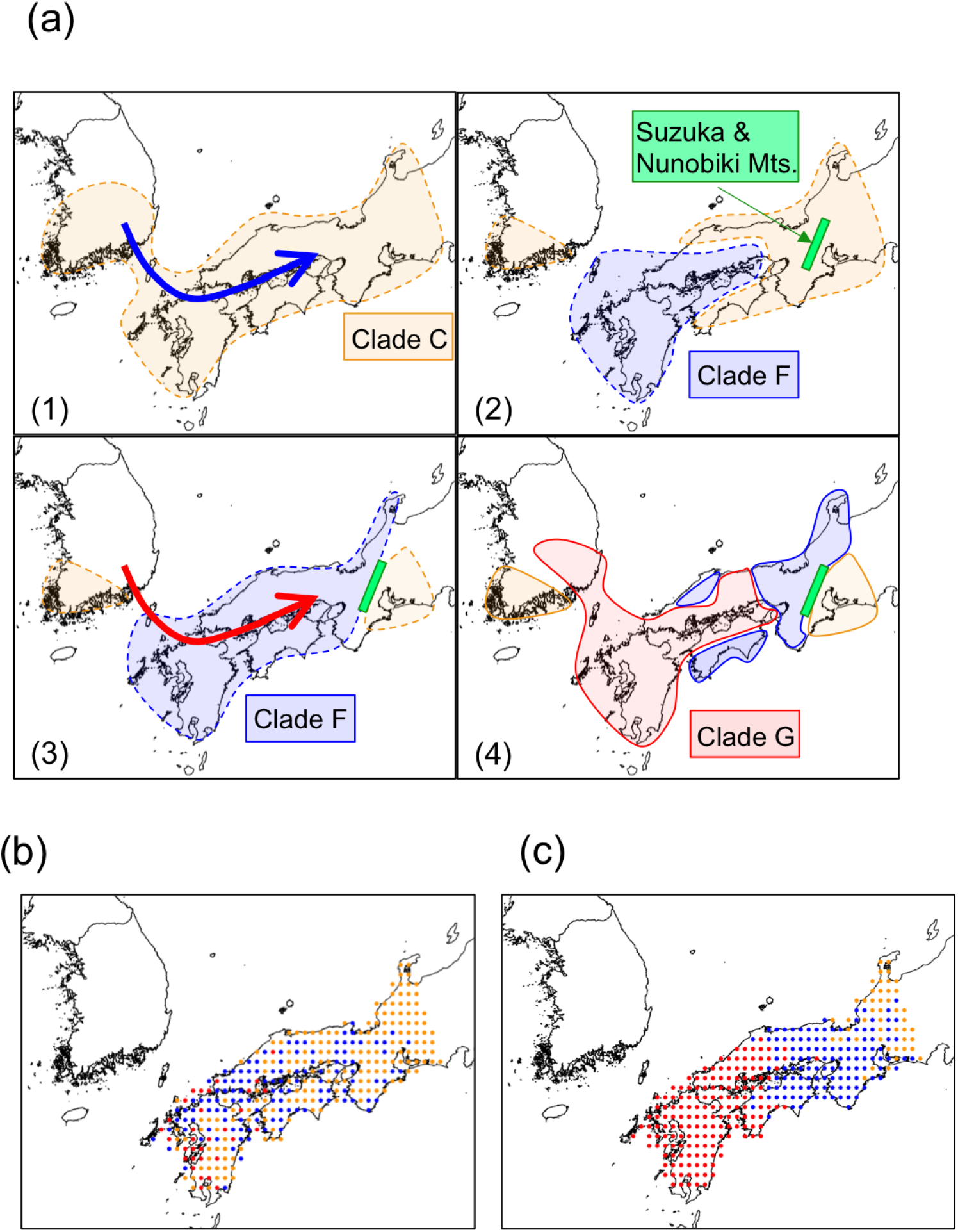
Schematic view and scenario of the formation process. (a) scenario with replacement for *Nipponocypris temminckii:* The number (1–4) at the bottom of each window represents the sequence in time and migration. The uplift of Suzuka and Nunobiki Mountains occurred in this process (2). (b) and (c) are examples of simulated *N. temminckii* distributions. (b) is the neutral scenario (*m* = 0.5, *s* = 20.2, α = 0.5, *r* = 0.853), and (c) is the best scenario with replacements (*m* = 0.345, *s* = 20.2, α = 0.774, *r* = 0.853). In the neutral scenario, the distribution of new migrants would expand where native populations had existed, and several clades would mix in a wide area.

The estimated migration time of clade G is mostly consistent with the Bayesian time estimation under a relaxed clock. The Bayesian time estimation by BEAST dates the timing of migration of clade G as 1.120 Ma [0.713-1.430]. As the estimated timing of migration of clade F is 1.31 Ma, clade G migrated to Japan 1.013 Ma [0.696–1.263].

Clade F is widely distributed in western Japan, but the distribution is isolated by Clade G in Chugoku and Shikoku. This pattern suggests that migration of clade G occurred after migration of clade F had settled into this habitat. Based on the fossil record, it is assumed that a land bridge between Kyushu and the continent of Asia formed at 0.43 Ma, 0.63 Ma, 1.2 Ma, and at about 5.3 Ma (Taruno, 2010). At the Last Glacial Maximum, sea-level was about 120 m lower than present (Rohling et al., 1998). Since the current minimum water depth between Korea and Kyushu is 130 m, the glacial periods were not necessarily accompanied by land-bridge formation. Furthermore, not all migration events contributed to the endemicity of Japanese land mammals (Sato, 2017). While the migration of clade F and subsequent migration of clade G occurred in the Pleistocene, the exact timing of migration remains unresolved.

### 3.4. *T. limbata* and related species, and *C*. spp. and *C. cuvieri*

Three clades (1, 3, and 4) of *T. limbata* were consistently identified in western Japan (Figure 6a). Sister clades (5–7) occurred in Korea, where they were classified as one of *T. koreensis, T. latimarginata* Kim, Jeon & Suk, 2014 or Korean bittering *T. signifer* (Berg, 1907). Clade 2 includes *T. limbata* from a mountainous location of Japanese Chugoku, and *T. somjinensis* from the upper reaches of the Korean Seomjin River. The fact that clades 1–4 are monophyletic, and the distribution of *T. somjinensis* is restricted to the upper Seomjin River where it is surrounded by *T. koreensis* and *T. latimarginata* (Jeon et al., 2018), suggests the *T. limbata* lineage formerly had a continuous distribution from Korea to Japan, but that this has since been divided by *T. koreensis* and *T latimarginata*. The divided distributions of clade 2 also imply its past continuous range from Korea to Chugoku in Japan, with clades 3 and 4 occurring in between.

**FIGURE 6.**
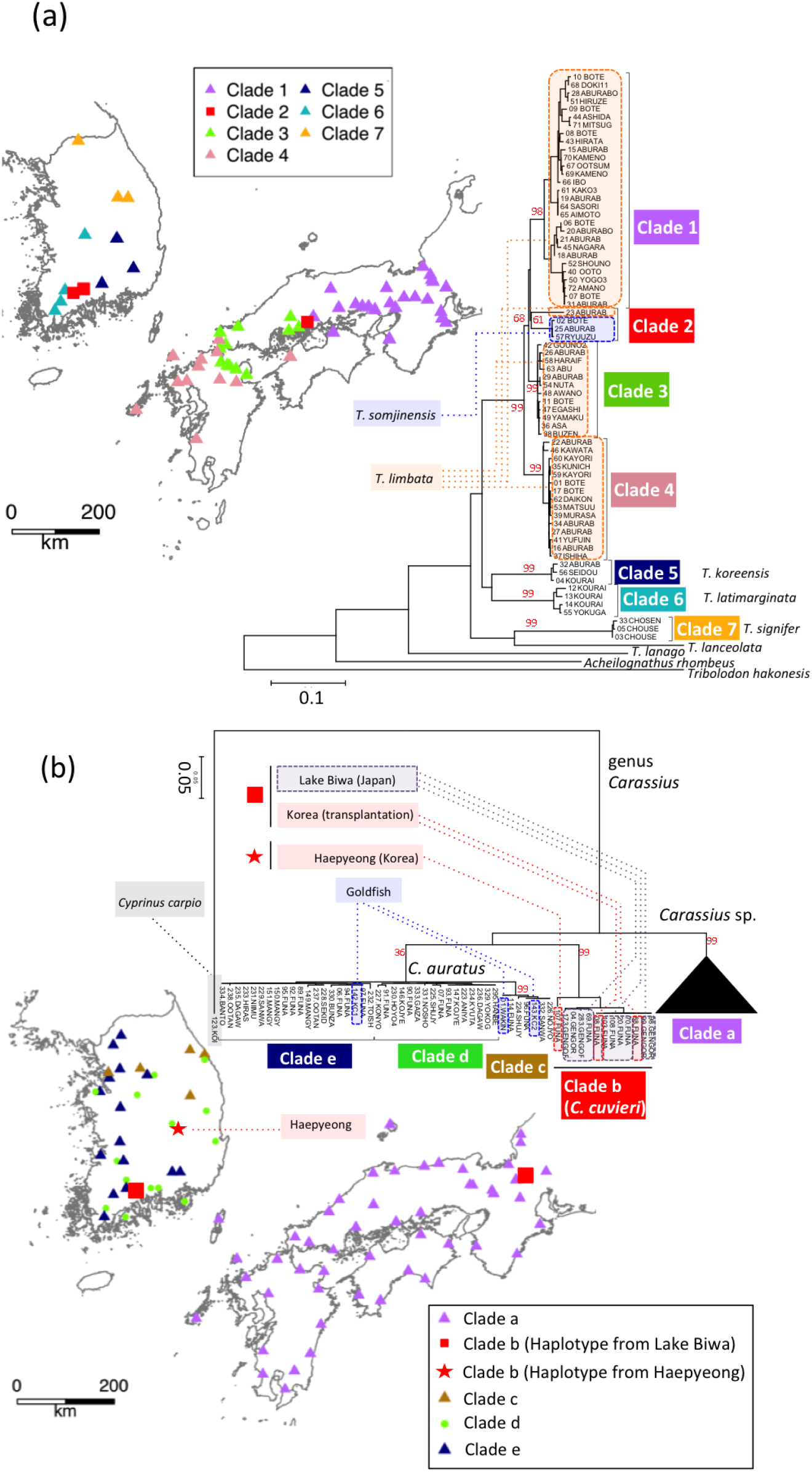
Phylogenetic trees of ND2 sequences and geographic locations: (a) *Tanakia limbata* and related species, (b) *Carassius* spp. As for *Carassius cuvieri*, the haplotype from Lake Biwa was found at a southern site in Korea (red rectangle). The haplotype from Haepyeong (red star) was very close to the haplotypes from Lake Biwa, but differed morphologically (it was initially identified as an individual from Korean clades c, d, and e).

*Carassius cuvieri* is endemic to Lake Biwa, Japan, and is represented by clade b in Figure 6b. Since *C. cuvieri* was artificially transplanted to various places in Korea and Japan, the same sequences as the original population in Lake Biwa were found in Korea. However, in the Korean region of Haepyeong we detected another sequence, which we presume is indigenous unless the same sequence occurs in Lake Biwa; the fish differs morphologically and had initially been identified as an individual from Korean clades. As transplanted *C. cuvieri* are morphologically indiscernible from *C. cuvieri*, it is unlikely that these Haepyeong individuals were transplanted. It is possible that clade b was previously continuously distributed in western Japan and Korea and that it has since become isolated by other clades in Korea.

## 4. Discussion

The Japanese freshwater fish populations we examined were derived from one or few clades of Korean populations. Other vertebrates in Japan such as the Siberian weasel *Mustela itatsi* Pallas, 1773 and the Japanese tree frog *Hyla japonica* (Günther, 1859) derived from one of a few clades of the continent as well. These phenomena suggest that the geographical origin of Japanese taxa is from Korea, and the migration waves out of Korea should be key factors in their distribution patterns if the Japanese taxa have related ones in Korea. The effects of migration might differ among taxa. In the case of *H. longirostris*, migration occurred only once, and the homogeneous genetic structure of this species in Japan indicates that dispersal occurred over a short period of time. The distribution of *H. longirostris* expands around the Inland Sea. To the contrary, *N. temminckii* migrated into Japan several times, probably resulting in the spatial structure of clades in Japan. The intermittent formation of the land bridge and stochastic success of migration through the land bridge generated taxon-specific waves into Japanese archipelago.

Genetic data from samples of multiple freshwater fishes from the Korean Peninsula and Japanese archipelago suggests waves of migrations of species from Korea have established themselves in Japan. Previous studies have interpreted the genetic structure of Japanese freshwater fish to have been caused mainly by vicariance. For example, the uplift of Suzuka and Nunobiki Mountains in the early Pleistocene (Figure 4c; Biwako Shizenshi Kenkyukai, 1994) caused genetic differences in fishes between the Tokai and Kinki regions (Tominaga et al., 2020). However, by incorporating phylogeographic data from Korea, we demonstrate that genetic differences in *N. temminckii* across Suzuka and Nunobiki Mountains are derived from different migration events from Korea, with the divergence of the two clades dating to the Pliocene (6.15 Ma; Figure B2) being significantly older than the uplift of the mountains. We also found older clades to be discontinuously distributed, separated by newer migrants. Such distributions cannot be fully explained by processes of diffusion or vicariance (Ronquist and Sanmartín, 2011), which predict genetic differentiation among components of division (Figure 1). If geographically divided individuals of older clades are genetically similar, their distribution was formerly continuous until recently because gene flow is extremely low in freshwater fishes. Extinction events and the expansion of distributions of other clades produced existing patterns.

Our hypothesis, that intra- and inter-species replacement has occurred in the process of successive migrations of taxa from Korea, explains the discontinuous distributions of freshwater fish taxa in Japan. The simulation suggests that the effects of replacements are significant compared to the null model of neutrality. In addition, divisions between clades exist in areas where dispersal barriers exist. For *N. temminckii*, the distribution boundaries occur around uplifted mountains (Suzuka and Nunobiki Mountains; Figure 4c) and the east-west axis of mountains in the Chugoku and Shikoku regions (Figure 4c, 4d). East of the Inland Sea, the uplifted Suzuka and Nunobiki Mountains presumably acted as barriers to dispersal for newer migrants, in addition to preventing replacement of an older clade (Clade C) in the Tokai region. A few samples of *H. longirostris* from the Tokai region probably exist because of artificial transplantation (Higuchi, 1980), as sequences from these individuals are identical with samples from the Kinki region. Therefore, the eastward dispersal of *H. longirostris* was stopped in the same way as it was for *N. temminckii*. The most recent migrant (clade G) had a continuous distribution around the Inland Sea, with an older clade (F) found in rivers discharging into the Sea of Japan, the Pacific Ocean, and in the upper parts of rivers flowing into the Inland Sea. Dispersal of the newer migrant probably caused replacement, with the old clade divided into an area where the former could not reach.

The formation of refugia might also contribute to discontinuous distributions. Many terrestrial animals in Japan sought refugia during glacial periods by migrating south or to low-altitude areas, and expanded their distributions again during interglacial periods (Sato, 2017). However, dispersal opportunities between rivers for freshwater fishes are more limited; they can either overwinter in local springs (Power et al., 1999) or go extinct. Therefore, we postulate that the main factors contributing to the present-day distributions of Japanese freshwater fishes are not the distributions of climatic refugia, but divergence, dispersal (Kitagawa et al., 2003; Takehana et al., 2003), migration and replacements.

The signature of replacements is also found in other freshwater fishes in Japan. For example, Japanese rice fish *Oryzias latipes* (Temminck & Schlegel, 1846) in western Japan has a divided distribution (Iguchi et al., 2018; Takehana et al., 2003), and the pike gudgeon *Pseudogobio esocinus* (Temminck & Schlegel, 1846) (Watanabe et al., 2017) and Japanese spined loach *Cobitis biwae* Jordan & Snyder, 1901 (Kitagawa et al., 2003) have older clades in eastern and parts of western Japan, and more recent clades in wide areas of western Japan. Our hypothesis may not be restricted to freshwater fishes, as the distributions of closely related moles, *Mogera wogura* (Temminck, 1842) and *M. imaizumii* (Kuroda, 1957), are also similar to that reported for *N. temminckii*, with *M. wogura* expanding from Korea to western Japan and *M. imaizumii* in eastern Japan and some isolated area in western Japan (Abe, 1995, 2001; Figure B5). These distributions are parapatric, with the latter having been replaced by the former at the distribution boundary; replacement of *M. imaizumii* by *M. wogura* was considered a decisive factor in the formation process (Abe, 2010).

An interesting problem is whether replacements are confined to mitochondrial genomes, or if they extended to nuclear genomes also. Allozyme analysis of *N. temminckii PEPA* nuclear locus revealed two-allele polymorphism at the upper portions of a few rivers located in the western Tokai region. In the Tokai region where clade C was fixed, the allozyme haplotype was fixed (*120) except the above populations. In the Kinki region where clade F was fixed, the allele is fixed to another (*100). The observed number of individuals by genotype are consistent with the Hardy-Weinberg equilibrium. The haplotypes of mitochondria and allozyme were not consistent, which means that the individuals over the boundary crossed randomly. They were probably caused by stream capture (Table A1 in Appendix A). Since clades F and G are genetically much closer than clades C and F, they could also hybridize. This implies that replacements occurred through change in composition of admixture. In the case of rosy bitterling *Rhodeus ocellatus* (Kner, 1866), when a non-native subspecies from the continent of Asia (*R. o. ocellatus*) was introduced into a population of Japanese native freshwater *R. o. kurumeus*, the mitochondrial and nuclear DNA of the former was replaced by the latter through hybridization (Kawamura et al., 2001). According to Ohta (1972), the efficiency of selection is negatively correlated with population size. The Korean Peninsula is part of the continent of Asia, and continental populations might have experienced higher competition than Japanese ones, and accordingly have higher fitness. Since newer migrants might have experienced higher selection pressures on the continent for a longer time, they may have higher fitness than indigenous species, resulting in the older clade being replaced by the newer clade. Japanese giant salamanders *Andrias japonicus* (Temminck, 1836) (Nishikawa, 2017) and Japanese weasel *Mustela itatsi* Temminck, 1844 (Imaizumi, 1960) are being replaced by related species that have been artificially transplanted from Korea or the continent of Asia.

Intra- and inter-species replacements have also occurred in Korea, where Korean *C. cuvieri* at Haepyeong occurred upstream of Nakdong River, surrounded by the sister clade, *C. auratus*. This pattern is similar to the Korean *T. somjinensis*, which comprises a monophyletic clade (Clade 2) with Japanese *T. limbata*, and inhabits the upstream waters of Seomjin River, surrounded by *T. koreensis* and *T. latimarginata*. Local distribution barriers such as waterfalls or flashy streams may have helped these isolated clades survive, or survival may be due to chance. In southwestern Korea, we sampled populations of *H. longirostris* (Clade II), comprising a monophyletic clade with the Japanese clade, which were geographically surrounded by other clades. In the case of *N. temminckii*, the clade from southwestern Korea (Clade C) also had a patchy distribution around this region. The Japanese tree frog *H. japonica* (Figure B6) and Siberian weasel *M. itatsi* (Shalabi et al., 2017; Figure B7a) have divided distributions of certain clades on Tsushima Island, a small island between Korea and Japan in the middle of the migration route, with populations in the middle of the Korean Peninsula or Russia comprising monophyletic clades with those on Tsushima. Other clades were sampled in between (e.g., southern Korea). In these two species, we expect that the latter clades replaced the former clades in Korea, but the latter could not migrate to Tsushima Island. Both *T. somjinensis* and *N. koreanus* are regarded as Korean endemic species, but they are components of monophyletic clades with different species in Japan. The taxonomy of these species requires review.

While both replacements and waves of migration out of Korea are essential factors in phylogeography, other factors need considering also. For example, *C. cuvieri* migrated into Japan in a single wave because its population is monophyletic. However, the distribution of this species is restricted to the freshwater system of Lake Biwa; as it prefers calm waters, the loss of paleo-river systems around the Inland Sea with rising sea level may have reduced suitable habitat. A similar pattern was detected for the freshwater fish three-lips *Opsariichthys uncirostris* (Temminck & Schlegel, 1846) and related species (Okazaki et al., 2002). The population of *T. limbata* in Japan was segregated into 4 clades. Among them, only single clade (clade 2) contained the Korean population. To better understand the phylogeographic structure, a more comprehensive model is necessary. We analyzed one mtDNA locus and one nuclear DNA locus. Genome data would provide more information to assist with understanding the history of these taxa in Korea and Japan. Nevertheless, our hypothetical process involving waves of migrations and replacements can be applied for other places and for different taxa, such as the brown bear *Ursus arctos* (Hirata et al., 2013, 2014; Waits et al., 1998; Figure B7b) and divided distribution of the mountain hare *Lepus timidus* Linnaeus, 1758 (Kinoshita et al., 2012) in Hokkaido. Chinese rice fish *Oryzias sinensis* Chen, Uwa & Chu, 1989 had a divided distribution in Korea, which might be replaced by a clade migrating from the Continent of Asia (Takehana et al., 2004; Figure B8). Although our findings owe much to the suitable geographical conditions of the Japanese archipelago and Korean Peninsula, waves of migrations and replacements may be more common and have more widely influenced the formation of biota than previously recognized.

## Acknowledgements

We deeply thank Jeffrey L. Thorne for valuable comments and suggestions on analytical methods and Tomoaki Hori for technical advice in speeding up the simulation program. We also thank Steve O’Shea (PhD) from Edanz Group (www.edanzediting.com/ac) for editing a draft of this manuscript. This study was supported by Grants-in-Aid for Scientific Research (B) 16H02788 and 19H04070 from the Japan Society for the Promotion of Science.

## Supporting information

Additional supporting information may be found online at the end of this article.

## CRediT authorship contributions statement

Shoji Taniguchi: Conceptualization, Methodology, Data Curation, Writing—Original Draft, Visualization.

Johanna Bertl: Methodology, Writing—Review & Editing.

Andreas Futschik: Conceptualization, Methodology, Writing—Review & Editing. Hirohisa Kishino: Conceptualization, Methodology, Writing—Review & Editing, Project administration, Funding acquisition.

Toshio Okazaki: Conceptualization, Investigation, Resources, Data Curation, Writing— Review & Editing, Project administration.

