## Supporting information for "Waves out of the Korean Peninsula and inter- and intra-species replacements in freshwater fishes in Japan"

**S1 Appendix**

1. **PCR and sequencing of ND II and cytochrome b**
   1.1 **PCR**

*Hemibarbus longirostris*: Following Hall & Nawrocki (1995), we carried out PCR on the ND1-16SRNA region of mtDNA (about 2.0 Kbp) using the following primers.

Forward: 5′-ACCCCGCCTGTTTACCAAAAACAT-3′

Reverse: 5′-GGTATGAGCCCGATAGCTTA-3′

Fifteen types of restriction enzymes were utilized: AciI, AfaI, AluI, BfaI, BstUI, DdeI, HaeIII, HhaI, HinfI, MboI, MspI, NlaIII, ScrFI, Sau96I, and TaqI.

*Nipponocypris temminckii*: the same primer as that for *H. longirostris* was used. Thirteen types of restriction enzymes were utilized: AfaI, AluI, BstUI, DdeI, HaeIII, HhaI, HinfI, MboI, MspI, NlaIII, ScrFI, Sau96I, and TaqI. From the cleavage type of each enzyme and the results of sequence analysis, it was found that individuals in clades F and G can be distinguished by differences in the cleavage type of BstUI, DdeI and TaqI.

*Carassius* spp.: the same primer as that for *H. longirostris* was used. Ten types of restriction enzymes were used: AfaI, BfaI, BstUI, HhaI, HinfI, MboI, MspI, NlaIII, ScrFI, and TaqI were adopted. The cleavage patterns of six enzymes could discriminate *C. cuvieri* from other *Carassius* species.

*Tanakia limbata*, *T. koreensis* and its related species: following Palumbi et al. (1991), we performed PCR on about 2.2 Kbp, including the control-12 SRNA regions of mtDNA, using the following primers.

Forward: Cb3R-L: 5′-CATATTAAACCCGAATGATATTT-3′

Reverse: 12SAR-H: 5′-ATAGTGGGGTATCTAATCCCAGTT-3′

Fifteen types of restriction enzymes were utilized: AciI, AfaI, AluI, BfaI, BstUI, DdeI, HaeIII, HhaI, HinfI, MboI, MspI, NlaIII, ScrFI, Sau96I, and TaqI.

**1.2 Sequencing**

First, total DNA was extracted from white muscle tissue (Asahida, Kobayashi, Taitoh, & Nakayama, 1996). Next, a partial region of the mitochondrial gene ND II was amplified by PCR using the following primer pairs designed based on the mtDNA sequence of *Cyprinus carpio* (Chang, Huang, & Lo, 1994): (5′-TWTYGGGCCCATACCCCRAA-3′) and (5′-GCTTTGAAGGCTYTTRGTCT-3′). PCR was conducted for 30 cycles at 94°C for 1 min, 52°C for 1 min, and 72°C for 2 min. The amplified DNA product was purified with a QIA quick PCR Purification Kit (Qiagen, Germany), and sequences were determined by an automated DNA sequencer (Applied Biosystem 377A). Cytochrome b sequence data were obtained in the same way. Primers included: L14724 (Palumbi et al., 1991) (5′-TGACTTGAARAACCAYCGYYG-3′) and H15915 (Aoyama, Watanabe, Ishikawa, Nishida, & Tsukamoto, 2000) (5′- ACCTCCGATCTYCGGATTACAAGAC-3′).

2. **Divergence time estimation**

The topology of the Maximum Likelihood (ML) tree and sequence data were imported into MCMCTREE package. In MCMCTREE analysis, we assumed HKY85 as the model of nucleotide substitution and the *correlated rate* as the model of evolutionary rate. Since no reliable fossil record was available to set calibration points, we estimated relative age by setting the time at the tree root as 1. Additionally we defined a few loose calibrations based on ML and Bayesian trees (Figure B2a, B2b) to prevent poor mixing of MCMC due to undue deviation from the reconstructed phylogenetic trees, which would not influence the order of node 1, 2 or 3. That is, we set four calibrations (Figure B3), node 4: ' < 0.5 ', node 5: ' < 0.8 ', node 6: ' < 0.75 ', and node 7: ' > 0.99 < '. Default values were adopted for other hyper-parameters. We performed MCMC simulation, and sampled the parameter set every 2000 iterations. By excluding the first 2000 parameter sets as burn-in, we obtained the final MCMC sample of 20,000 parameter sets. We estimated the posterior distribution of the difference between the relative ancestral age of clade C and F, as well as clade C and G, by calculating differences in the MCMC sample. Given the migration of clade F as 1.31 Ma, the conditional posterior of the migration period of clade G was estimated from MCMC samples obtained with BEAST and MCMCTREE. We calculated the ratio between migration times of clades G and F for each sample and then multiplied them by 1.31.

**3. Simulated distribution-formation process of *Nipponocypris temminckii***

We simulated the formation process of biogeographic distribution by generating the dynamics of the states on the 329 evenly distributed lattice-like grids. Their envelope covered the whole range in distribution of *N. temminckii* throughout Japan, expanded to the estimated coastal line at the time of glaciation periods (120 m below the current sea level; Figure 2) (Fairbanks, 1989; Rohling et al., 1998). The distance between points was defined as the geographic distance. Distances were calculated using the R (R Core Team, 2017) package geosphere (Karney, 2013).

As reported in our Results, phylogeographic analysis implied a specific formation scenario. That is, clade C arrived first, and clades F and G in turn migrated into Japan from the Korean Peninsula. We assumed that clade F had migrated into western Japan 1.31 Ma (Figure B2b). The state of the simulation was the clade assignment of each grid. The simulation started with the state of assignment to clade C except for the three points at northern Kyushu, which were assigned to clade F. In *r* Ma, clade G migrated. At each simulation step, the clade at each point had multiple offspring. One stayed at the same point and the others dispersed to nearby points. We assumed the distance (*x*) that an offspring dispersed within time *t* followed a gamma distribution, in accordance with:

$Gamma\left( shape={t\cdot m}/s,scale=s \right)$	(1)

where *m* is the expected distance that an offspring disperses per unit time (km/Kyr), and *s* is the scale parameter. In this model, the mean dispersal distance in time *t* is *tm*, and the variance is *tm∙s*. The probability that an offspring disperses from point *i* to point *j* is

$p_{ij}=P\left( X>d_{ij} \right)$	(2)

where *d_ij_* is the distance between points *i* and *j*. If more than one clade coexists at point *i* as a result of dispersal, one clade was chosen randomly with a probability that reflects the difference in the fitness between clades. As a simple model representing the fitness difference between clades, we assumed that both the selective advantage of clade F over clade C, and that of clade G over clade F, was *α*, and that the selective advantage of clade G over clade C, was *α^2^*. With this model, the replacement probability in a case of clade-coexistence is:

$p\left( F | C,F \right)=p\left( G | F,G \right)=\alpha$,

$p\left( C | C,F \right)=p\left( F | F,G \right)=1-\alpha$,

$p\left( G | C,G \right)={\alpha^{2}}/\left\{ \alpha^{2}+\left( 1-\alpha\right)^{2} \right\}$,

$p\left( C | C,G \right)={\left( 1-\alpha\right)^{2}}/\left\{ \alpha^{2}+\left( 1-\alpha\right)^{2} \right\}$,	(3)

$p\left( C | C,F,G \right)={\left( 1-\alpha\right)^{2}}/\left\{ \alpha^{2}+\alpha\left( 1-\alpha\right)+\left( 1-\alpha\right)^{2} \right\}$,

$p\left( F | C,F,G \right)={\alpha\left( 1-\alpha\right)}/\left\{ \alpha^{2}+\alpha\left( 1-\alpha\right)+\left( 1-\alpha\right)^{2} \right\}$,

$p\left( G | C,F,G \right)={\alpha^{2}}/\left\{ \alpha^{2}+\alpha\left( 1-\alpha\right)+\left( 1-\alpha\right)^{2} \right\}$.

**4. Parameter estimation** **by ABC**

**4-1 Fitting values of summary statistics to observed values**

The generated biogeographic distribution varied largely among runs of simulation. Instead of fitting the generated distribution by itself to the observed distribution, we fitted the value of summary statistics to the observed value. For each simulation run we calculated the values of summary statistics from the current states on the nearest grids to the sampling locations. These were contrasted with observed values.

To avoid excessive computational costs, the simulation comprised 40 discrete evenly spaced steps. Therefore, the parameter *r*, the timing of migration, was selected from 39 equally spaced values. Finally, with the function abc of the R (R Core Team, 2017) package abc (Csilléry, François, & Blum, 2012), the posterior distribution of each parameter was obtained using the neuralnet method (Blum & François, 2010) with a tolerance rate 0.025. In total 360,000 runs were conducted to estimate the posterior.

**4-2 Two Summary statistics**

A) Templeton statistics

Templeton proposed the clade distance (D_c_) and the nested clade distance (D_n_) in NCPA (Posada et al., 2006; Templeton et al., 1995). The clade distance measures the geographic spread of a clade, and the nested clade distance measures how a clade is geographically distributed relative to other clades (Figure B1). D_c_ and D_n_ were calculated as follows. From all the points in the distribution of *N. temminckii*, the centroid point of all the sampling points (C_All_) and the centroid point of clade X (C_X_) were extracted. The centroid point of clade X is the member $i\in cladeX$ that minimizes the average distance to the other members:

$argmin\sum_{j\in cladeX} d_{ij}$	(4)

where $d_{ij}$ is the geographic distance between points *i* and *j*. Then, D_c_ and D_n_ of clade X become:

$$D_{c}=mean\left( d_{C_{X}j} \right)$$

$D_{n}=d_{C_{X}C_{All}}$	(5)

B) Spatial autocorrelation

Spatial autocorrelation (S_a_) measures how each clade is aggregated or mixed (Figure B1). S_a_ is defined as

$S_{a}={\sum_{i,j} exp\left( -cd_{ij} \right)g_{ij}}/{\sum_{i,j} exp\left( -cd_{ij} \right)}$	(6)

Here, $g_{ij}$ is equal to 1 if individuals at points *i* and *j* are members of one clade, while it equals 0 if they belong to different clades. In this study, the value of *c* was set to 0.02. With this value, a point 10 km away has a weight of 0.82 and a point 100 km away has a weight of 0.14.

**4-3 Prior distribution**

Vague but informative priors of the four parameters were set as: $m\sim unif\left\{ 0,5 \right\},s\sim unif\left\{ 0,50 \right\}$*,* $\alpha\sim unif\{0.5,1\}$*,* $r\sim unif\left\{ 0,1.31 \right\}$*,* to avoid unduly poor convergence by sampling highly unlikely parameter values at high frequency.

**5 Allozyme analysis**

**Table S1**

**Genotypes and frequency at the *PEPA* locus and their clade type at the three river populations**

| Place | Genotype | Observed number  of individuals | Expected number  of individuals | Clade type |
| --- | --- | --- | --- | --- |
| Kumozu River | *100/*100 | 0 | 0.02 | - |
|  | *100/*120 | 1 | 0.95 | C |
|  | *120/*120 | 12 | 12.03 | - |
|  |  | X-squared = 0.0208 |  |  |
| Ibi River | *100/*100 | 1 | 1.13 | F |
|  | *100/*120 | 7 | 6.75 | F |
|  | *120/*120 | 10 | 10.13 | C |
|  |  | X-squared = 0.0247 |  |  |
| Suzuka River | *100/*100 | 2 | 1.07 | C |
|  | *100/*120 | 4 | 5.87 | F, F |
|  | *120/*120 | 9 | 8.06 | - |
|  |  | X-squared = 1.52 |  |  |

Allozyme analysis of the *PEPA* locus was conducted as described in Okazaki et al (1991). The upper portions of Kumozu, Ibi and Suzuka rivers located in the western Tokai region, polymorphism was observed caused by *100 and *120 alleles at *PEPA* locus. The observed number of individuals by genotype are consistent with the expected number by the Hardy-Weinberg equilibrium. Several individuals were sequenced from samples. Individuals with genotype *120/*120 were classified into clade C, and genotype *100/*100 into clades C and F. Heterozygous individuals were classified into clades C or F depending on the sample. The haplotypes of mitochondria and allozyme were not consistent, which means that the individuals over the boundary crossed randomly. We conducted a goodness-of-fit test by simulating the random values from the chi-square distribution (df = 1). We simulated the 3 random values at each iteration, chose the maximum value, and obtained the distribution of the maximum value. The observed maximum chi-square value was 1.52, which is far lower than the 95 percentile of the simulated distribution (5.65). Observed genotype frequencies are consistent with expected values.

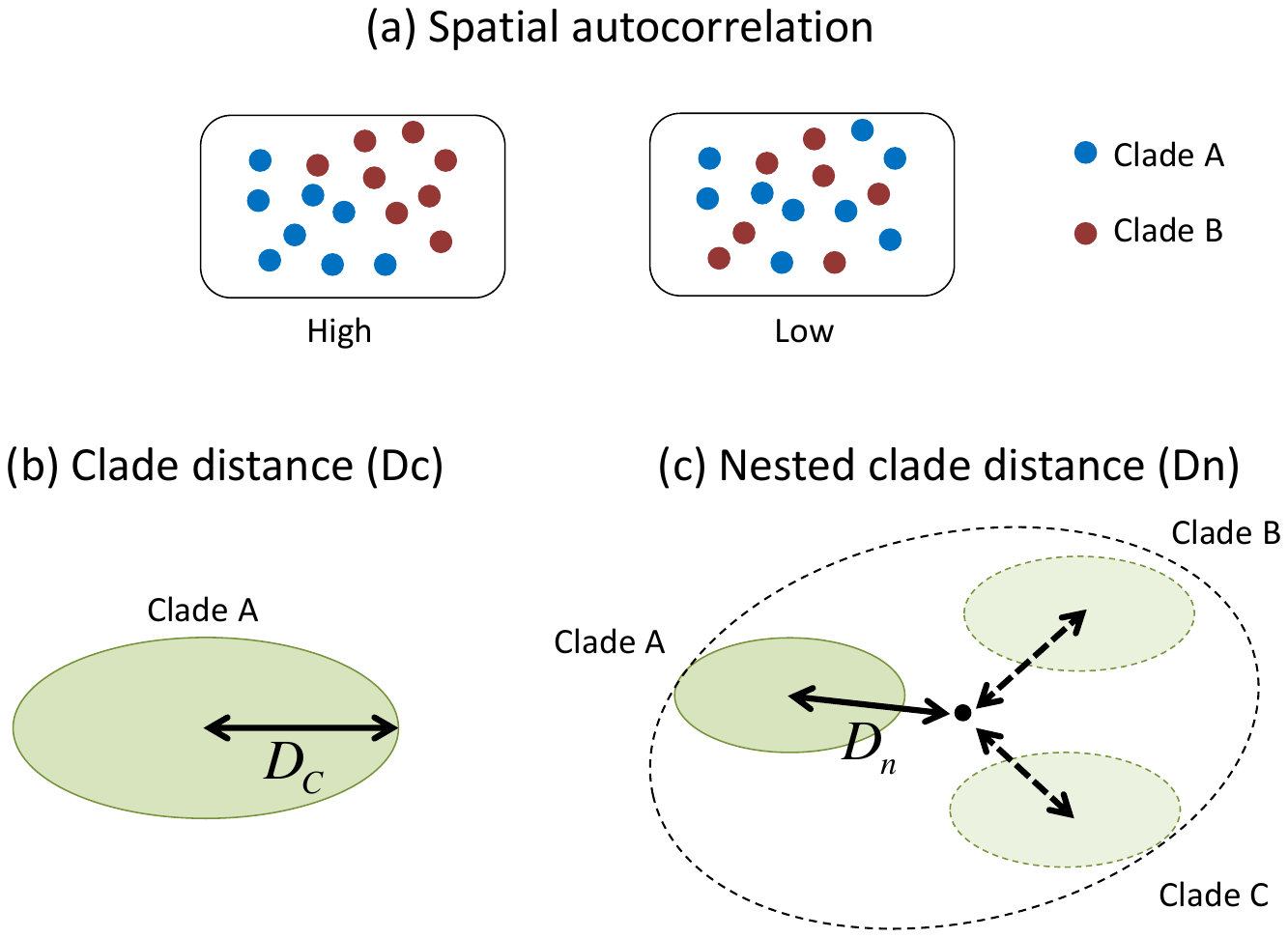

**Fig S1**

**Summary statistics for ABC-based testing hypothesis of intra-species replacement:** (a) spatial autocorrelation, measuring how each clade is aggregated or mixed; and (b) clade and (c) nested clade distances, measuring geographic arrangement and expansion of distribution.

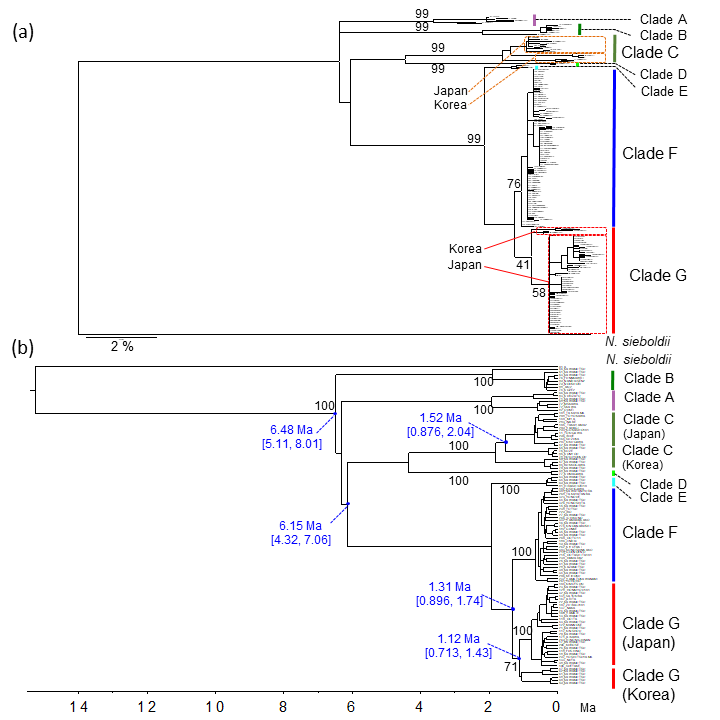

**Fig S2 Phylogenetic trees of *Nipponocypris temminckii* reconstructed from ND II sequences.** (a) ML tree, (b) Bayesian tree. Black numbers in phylogenetic trees indicate bootstrap or posterior probabilities (%). Blue numbers without brackets in (b) indicate point estimates of node age; blue numbers within brackets indicate 95% upper and lower credibility interval limits (Ma).

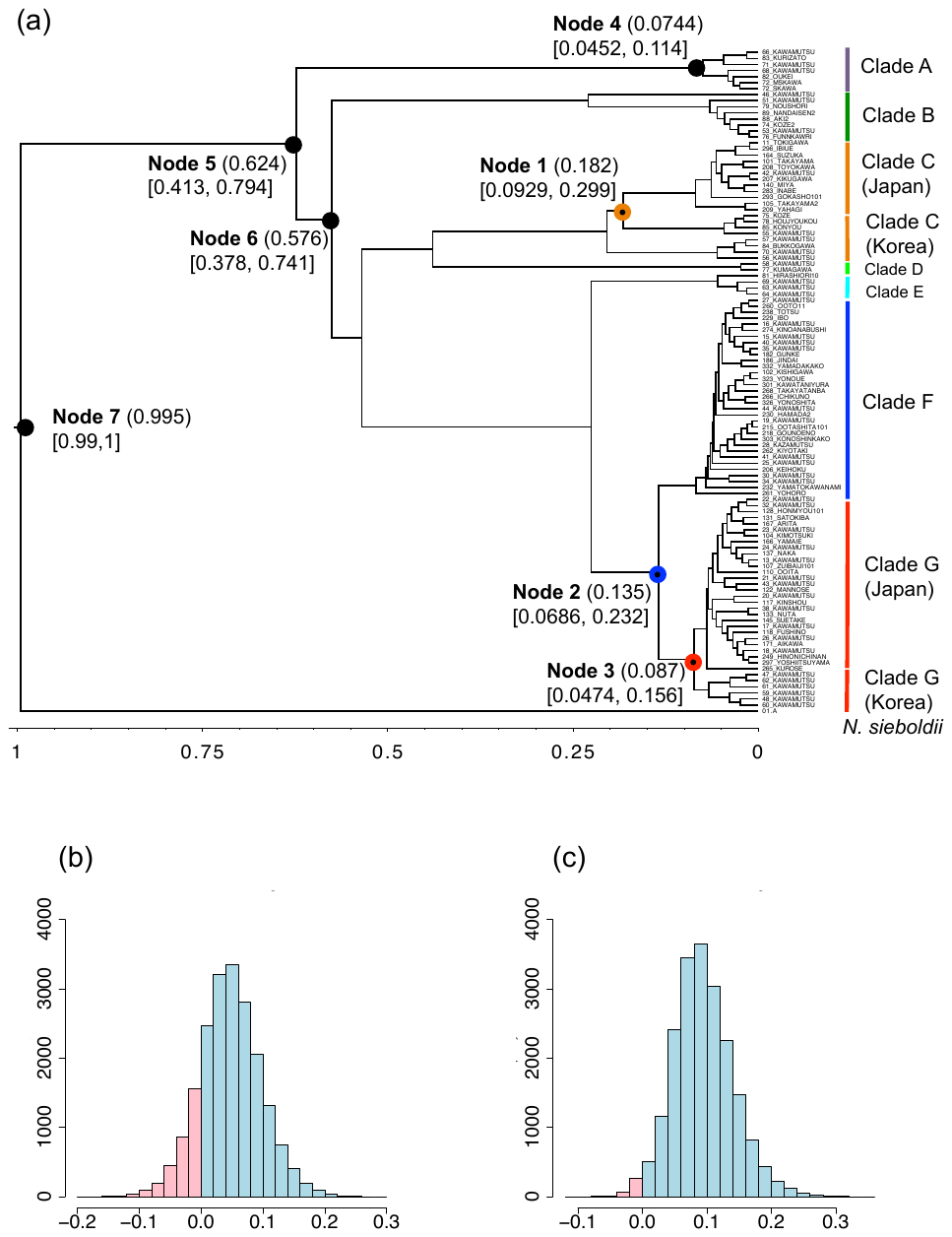

**Fig S3**

**Divergence time estimation by MCMCTREE:** (a) estimated relative divergence time of nodes in parentheses beside node number, and 95% credibility interval below the node number; (b) and (c) histograms of MCMC samples. (b) Difference in relative divergence time between nodes 1 and 2; the probability of node 1 being older than node 2 is 83.8% in blue, in contrast to the opposite event in red. (c) Difference in relative divergence time between nodes 1 and 3; the probability of node 1 being older than node 3 is 98.2%.

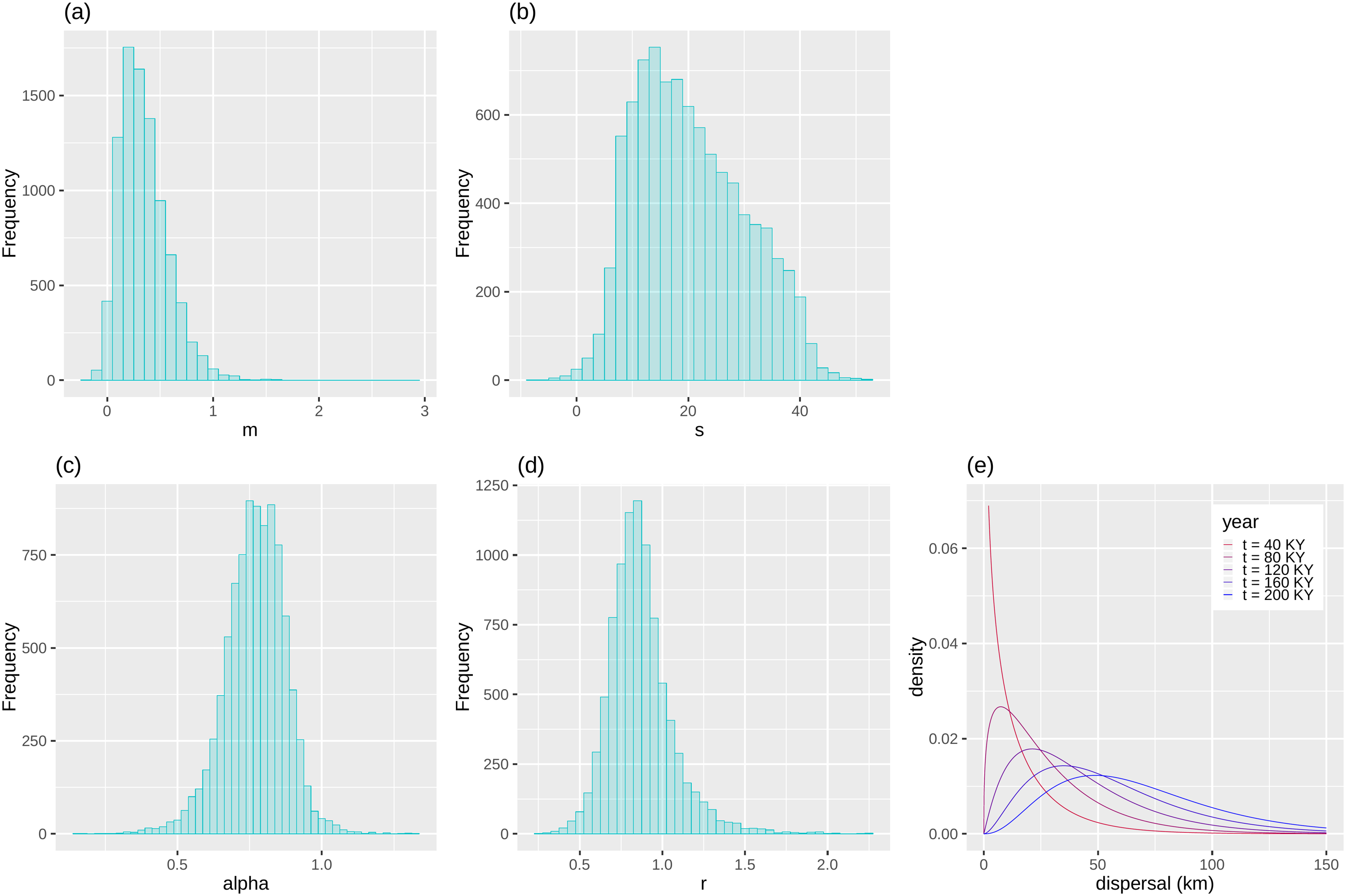

**Fig S4**

**Posterior distribution of four parameters, and probability density of dispersal distance:** Posterior distributions of (a) *m*, (b) *s*, (c) *α*, and (d) *r*. (e) The probability distribution of dispersal distance under the point estimates of parameter *m* (0.345) and *s* (20.2).

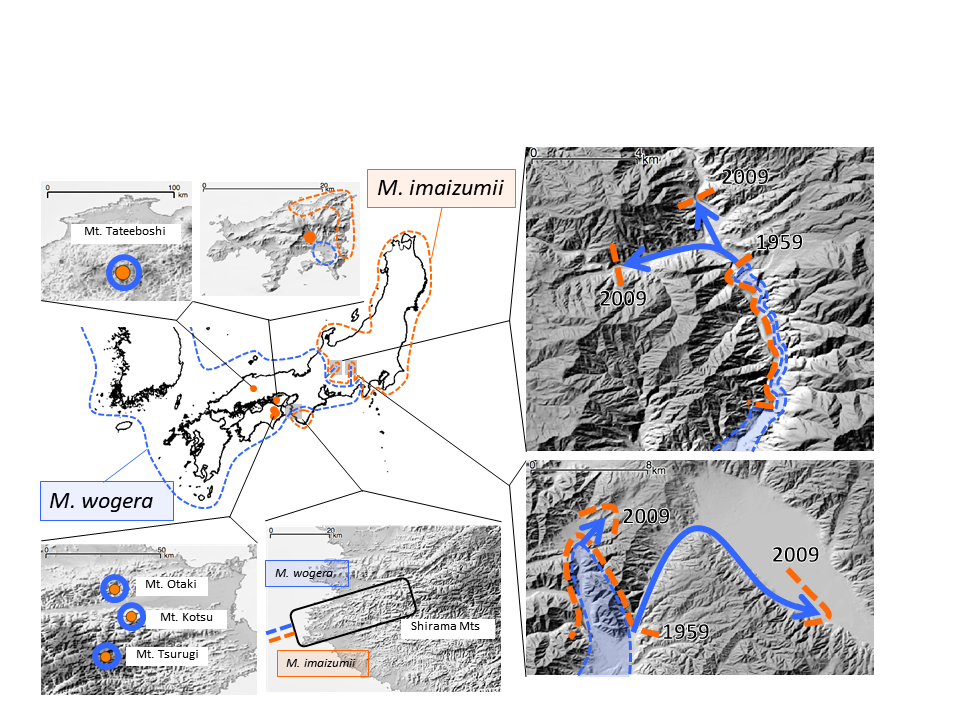

**Fig S5**

**Distribution of *Mogera wogura and M. imaizumii*.** At the eastern edge of the distribution of *M. wogura*, this species has reputedly expanded its distribution and replaced *M. imaizumii* between 1959 and 2009. In the Chugoku and Shikoku regions, the distribution of *M. imaizumii* is isolated to habitat atop several mountains or an island within the Seto Inland Sea; this species is also distributed in the southern Kinki region, with Shirama Mountains at the boundary. Map based on descriptions by Abe (1995, 2001, 2010). The original elevation chart (in color) was provided by the Geospatial Information Authority of Japan; marine areas were assembled using data from the Hydrographic and Oceanographic Department, Japan Coast Guard (Geospatial Information Authority of Japan, 2013).

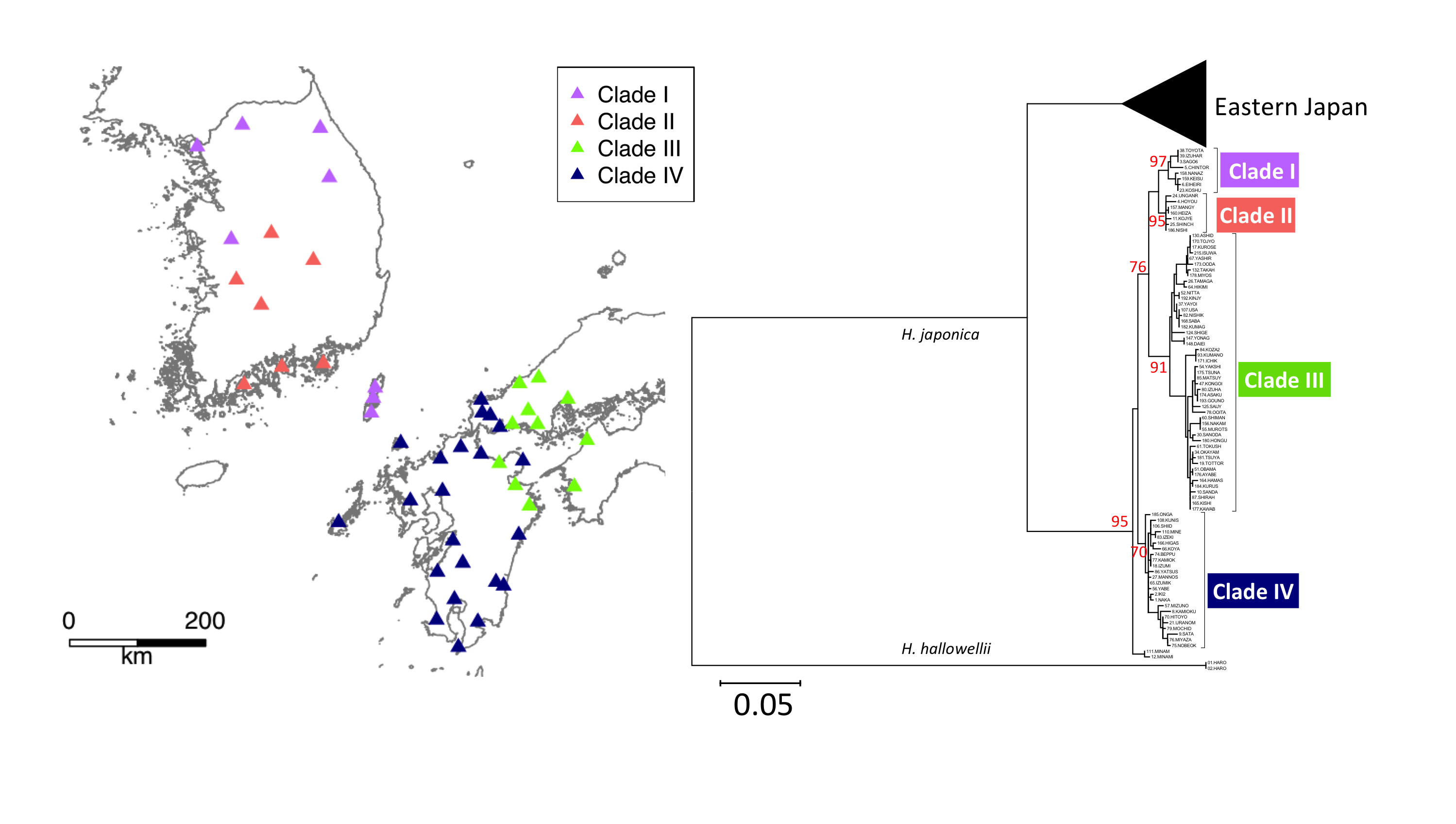

**Fig S6**

**Phylogenetic trees of ND II sequences and geographic locations: *Hyla japonica*.** ML phylogenetic trees and biogeographic maps for *H. japonica* were obtained in a similar fashion to those for *N. temminckii*. The red numbers were the bootstrap probabilities. We used sequences of *H. hallowellii* as an outgroup for *H. japonica*. Clade I was sampled in the middle of the Korean Peninsula and on Tsushima Island, Japan, while clade II was sampled in southern Korea. For *H. japonica*, the Korean Peninsula provided suitable habitat even during the last glacial maximum (Dufresnes et al., 2016), so it is unlikely that extinctions were caused by climate change. The distribution of clade I was divided by that of clade II. The average distance between *H. japonica* clades I and II is 1.87%, at the intra-species level. The obtained sequences were deposited in DDBJ/ENA/GenBank (accession numbers were LC568290- LC568534).

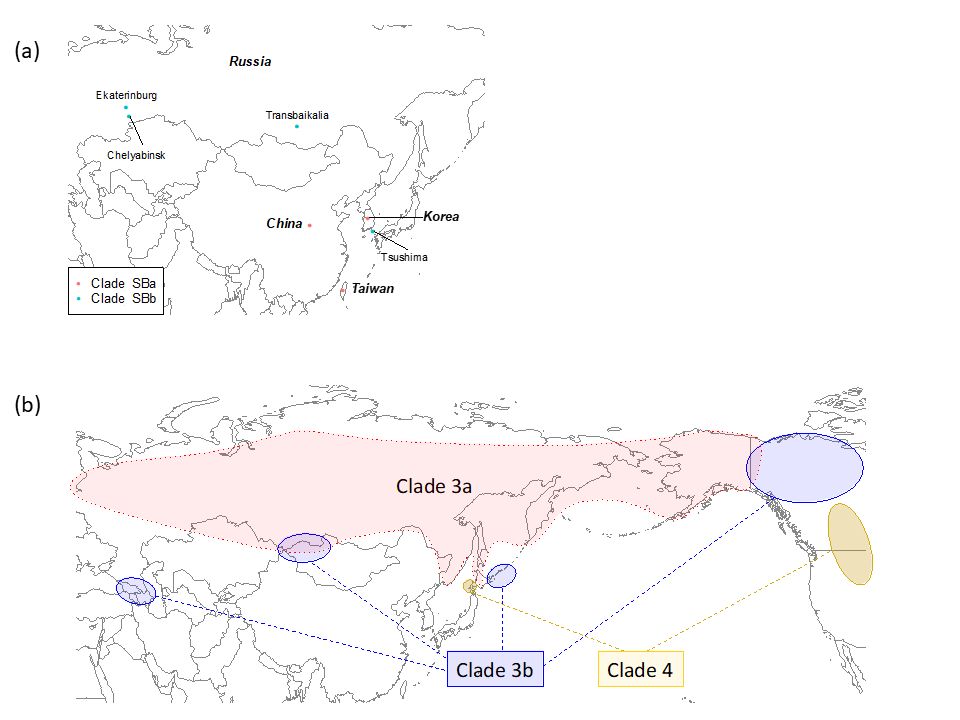

**Fig S7**

**Distribution maps of Siberian weasel (a) and Brown bear (b):** (a) modified from Shalabi et al. (2017). Clade SBa was sampled in China, Korea and Taiwan, while clade SBb was mostly sampled in Russia, but also Tsushima Island. The distribution of clade SBb is divided into Russia and Tsushima Island, while the distribution of clade SBa exists in between. (b) Based on previous studies (Hirata et al., 2013; Hirata et al., 2014; Waits et al., 1998) the distribution of clade 3a continuously expands from Russia to Alaska, while the distributions of other clades are divided. Clade 3b has several isolated populations on the Asian continent, around Japanese Hokkaido, and in North America. These isolated populations exist near the periphery of the continuous distribution of clade 3a. The distribution of clade 4 is also divided. Of the three clades, clade 4 in North America is farthest from Bering Strait; a population in the western area of Japanese Hokkaido exists also.

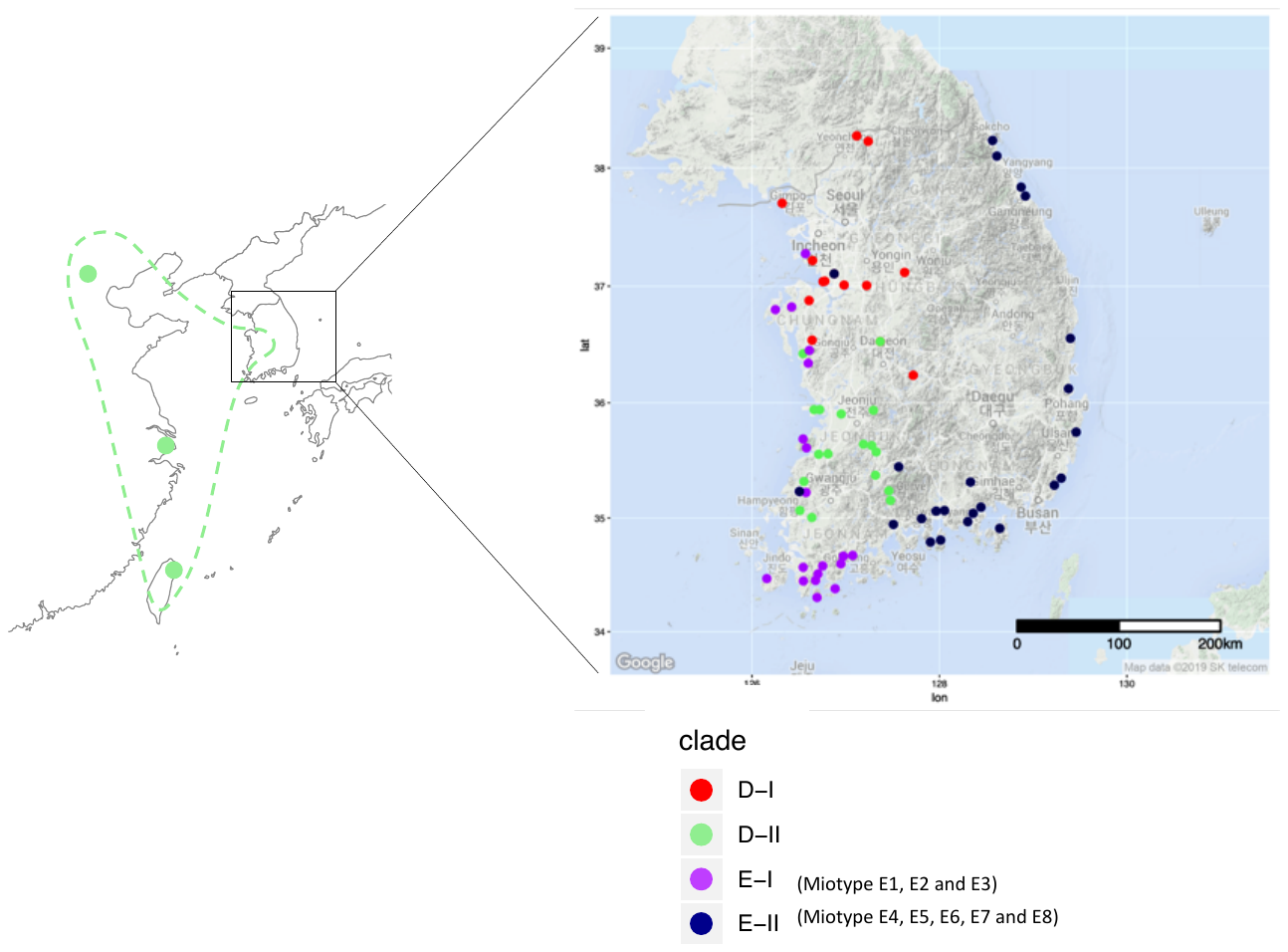

**Fig S8**

**Distribution maps for *Oryzias sinensis*, based on Takehana et al. (2004)**. Two clades (D, E) and two subclades of D (D-I and D-II) of *O. sinensis* are recognized; we also split clade E into subclades E-I and E-II. Subclade D-II has a distribution from China to southwestern Korea, and clade D-I inhabits midwestern Korea. Clade E-I occurs in the most southwestern part of Korea, with fragmented distributions where clade D-I and D-II are widely distributed. Clade E-II has a wide distribution in eastern Korea, and a fragmented distribution near Incheon and Hampyeong. Map drawn using the R (R Core Team, 2017) package ggmap (Kahle & Wickham, 2013).
